# Enhancing Retromer Complex Stability Ameliorates Synaptic Dysfunction in a Mouse Model of Alzheimer’s Disease

**DOI:** 10.1101/2024.06.11.598446

**Authors:** David Ramonet, Anna Daerr, Martin Hallbeck

## Abstract

Synaptic dysfunction is an early hallmark of Alzheimer’s disease, characterized by the disruption of synaptic transmission and plasticity. Central to these processes is endosomal trafficking, mediated by the retromer complex, which orchestrates the movement of vesicle contents for recycling to the plasma membrane, return to the Golgi, or degradation. Variants of VPS35, the cargo recognition component of the retromer complex, have been linked to neurodegenerative diseases, including Parkinson’s disease (PARK17, D620N mutation) and Alzheimer’s disease (L625P mutation). While substantial research has focused on Parkinson’s, the role of VPS35 in Alzheimer’s has been less explored. This study investigates the acute neuroprotective effects of retromer-stabilizing compounds in the 5xFAD mouse model of Alzheimer’s. Our results reveal that stabilization of the retromer complex not only mitigates pathogenic Aβ production mechanisms but also compensates for early synaptic dysfunction and microglial activation. Specifically, we observed significant modulation of genes involved in long-term potentiation and a reduction in abnormal retromer-associated cargos. These findings highlight the potential of retromer stabilisation as atherapeutic strategy to address fundamental pathological pathological processes in Alzheimer’s disease.

## Introduction

Synaptic dysfunction is a central feature not only in Alzheimer’s disease (AD) but across various neurodegenerative disorders. Defined as disruptions in synaptic transmission and plasticity, synaptic dysfunction is directly linked to cognitive decline and is recognized as one of the earliest hallmark of AD. The presence of beta-amyloid (Aβ) and tau proteins can directly disrupt neurotransmitter release, receptor signaling, and dendritic spine stability, thereby creating a causative link to synaptic anomalies (Bukke et al. 2020; Meftah and Gan 2023). However, the pathways underlying Aβ’s synaptic accumulation and potential intervention strategies remain incompletely understood, suggesting a complexity beyond initial assumptions.

Retromer complexes are critical for amyloid precursor protein (APP) and Aβ processing because they determine cargo selection for endosomal transport. Under normal physiological conditions, clathrin-mediated endocytosis initiates the formation of early endosomal vesicles, evolving into multivesicular bodies. These vesicles are subsequently routed back to the Golgi apparatus via the retrograde pathway or recycled to the plasma membrane. These vesicles contain a variety of receptors and also APP processing machinery. Thus, maintaining retromer complex integrity is vital for synaptic function (Small and Petsko 2020; Temkin et al. 2017; Walsh et al. 2021).

Central to the retromer complex is the VPS35 protein, which regulates the assembly and stability of the complex and is crucial for cargo recognition [Bonifacino2008]. Variants of VPS35, such as D620N, have been implicated in autosomal dominant forms of Parkinson’s disease (PARK17) (Vilariño-Güell et al. 2011; Zimprich et al. 2011), with other variants like L625P linked to early-onset Alzheimer’s disease (Rovelet-Lecrux et al. 2015). Early AD changes include modest reductions in VPS35 protein levels within the entorhinal cortex (Small et al. 2005).

Mutations in VPS35 disrupt its function, leading to APP accumulation in early endosomes, increased free Aβ and Aβ-exosome release, and enlarged axonal endosomes (Cataldo et al. 2000; Kim et al. 2016; Jiang et al. 2016). Furthermore, Aβ exposure has been shown to decrease VPS35 protein levels, with Aβ colocalizing with VPS35 (Ansell-Schultz et al. 2018) Interestingly, microglial dysfunction, significant in AD progression, is also modulated by the retromer complex, critical for appropriate microglial activation from the M0 state (Ren et al. 2022). Early AD changes include modest reductions in VPS35 levels within the entorhinal cortex (Small et al. 2005).

Two thiopenthene thioureas named R55 and R33, were identified during a screening aimed at stabilizing the VPS35-VPS29 complexation against thermal denaturation (Mecozzi et al. 2014). In vitro studies have demonstrated that these pharmacological chaperones, at low micromolar concentrations (5 μM), enhance VPS35 and VPS26 protein levels in mouse primary cortical neurons and effectively reduce APP processing, thereby decreasing endogenous Aβ42 and Aβ40 production (Mecozzi et al. 2014). Despite these promising in vitro results, In vivo studies on the potential neuroprotection through adenoviral or pharmacological stabilization of VPS35, have been met with skepticism by the scientific community due to problems with scientific rigor (Vagnozzi et al. 2023; 2021; J.-G. Li, Chiu, and Praticò 2023).

However, it remains to be demostrated how much protection in the murine Alzheimer’s model is achieved by retromer stabilisation, and even less which pathways are involved in this neuroprotection, beyond the extrapolation from in vitro studies that Abeta processing is involved. The question is important because although the VPS35 interactome includes many key players in AD pathophysiology and is therefore well positioned to act as a putative critical failure hub, the VPS35 D620N clinical variants produce Parkinson’s and not Alzheimer’s (Wider et al. 2008), and the changes observed in VPS35 expression in both patients and mouse models are rather modest.

In this study we investigate the effect on signaling networks disrupted in the Alzheimer’s disease model 5xFAD/C57BL6, following acute intraparenchymal administration of the tool compounds R55 and R33. Our aim is to shed light on the downstream mechanisms affected by the stabilization of the retromer complex, thereby providing a detailed map of the molecular events and cascades triggered by these interventions. Given the existing gaps in our understanding of the pharmacokinetics of these molecular chaperones and challenges associated with long-term intraparenchymal delivery studies we focus on the immediate biochemical events triggered by these interventions.

## Materials and Methods

### Drugs

The thiophene thioureas R55 and R33 were sourced as hydrochloride salts from CaymanChem (Ann Arbor, MI) and MedKoo (Morrisville, NC), respectively. To prevent degradation caused by the susceptibility of the unprotected thiourea groups to hydrolysis and oxidation, we dissolved the salts in high-purity anhydrous DMSO to a concentration of 300 mM, aliquoted them, and stored them at - 70°C. Just minutes before brain microinjection, we thawed one aliquot and further diluted it to 10 mM under sterile conditions with freshly prepared 20% sufobutylether betacyclodextrin (Thermo Fisher) in saline and used it as a single dose. To ensure consistency throughout the study, a control vehicle containing the same proportions of DMSO and betacyclodextrin was prepared alongside the working solutions, which contained 3% DMSO. Although these tool compounds have not undergone proper pharmacokinetic evaluation, injecting 500 nL of these solutions into the cortex would result in an estimated 15 μM of compound, assuming uniform distribution throughout the brain. This concentration is three times the estimated Kd for VPS35 (5 μM) and is already saturated for retromer stabilization in primary neurons, as described previously (Mecozzi et al. 2014). The value is also approximately the estimated Ic50 for the Aβ reduction assay in primary neurons, as reported in the same study.

### Animals and surgeries

Sixteen-week-old female mice of the 5xFAD/C57BL6 strain, specifically B6, were used in this study. Cg-Tg(APPSwFlLon,PSEN1*M146L*L286V)6799Vas/Mmjax mice were obtained from The Jackson Laboratory (JAX) and locally mated with C57BL/6J from the same source. The compounds were stereotaxically administered to the lateral parietal associative cortex. The coordinates used were caudal 2.06 mm, lateral 1.5 mm and ventral 0.8 mm. The infusion was slow, at 100 nL/min, and the total volume was 500 nL. Twenty-four hours after the injections, the mice were humanely euthanized, and their brains were quickly dissected. For the transcriptomics study, a cube with a lateral dimension of 1 mm, centered on the injection point, was isolated from the cortex of each mouse, along with another identically sized and relatively positioned cube from the contralateral noninjected hemisphere. Care was taken during the microdissection of the cube to ensure that no parts of the corpus callosum were incorporated into it. The cubes were quickly submerged in RNAlater (Thermo Fisher) and frozen at -70°C. The assignment of the two compounds and the vehicle sets into groups of n=6 animals per condition was randomized. For each cage containing siblings, three sets were used. Personnel were blinded to the identities of the compounds and vehicle until the end of the data analysis. For the histology study, an independent set of animals (n=3 per condition) was subjected to an identical microsurgery protocol. However, they were perfused intracardially with PBS and 4% PFA in PBS before whole-brain dissection. All procedures and working protocols were approved by the animal ethical committee of Linköping University.

### Expression profiling

An RNeasy+ Mini Kit (Qiagen) was used to prepare total RNA for each sample with the assistance of a QIAcube automation system (Qiagen). Quality control was performed using Bioanalyzer RNA 6000 Nano Chips (Agilent), and all samples had an RNA integrity number (RIN) > 8.5. RNA was quantified with Quant-iT (Invitrogen) before constructing an indexed library with the Illumina Stranded mRNA Prep Kit. Deep RNA sequencing was performed using a P50 on a NextSeq 2000 (Illumina) with the standard Illumina pipeline. The paired reads were mapped to the mouse GRCmm39 genome (Jun 2020) with HISAT2, obtaining between 20 M and 40 M mappings for each sample. DESeq2 in R was used to conduct differential expression analysis comparing the injected hemisphere with the contralateral hemisphere, as well as the three different microinjections (vehicle, R55, and R33). The data processing and analysis were performed using R (R Core Team, 2023, www.r-project.org) and Bioconductor (Huber et al. 2015), utilizing the GO, InterPro, KEGG, Monarch, Pfam, SMART, STRINGdb, UniProtKB, and WikiPathways databases. Reference datasets were obtained from (Bliim et al. 2019) (GSE110908) and MODEL-AD (Model Organism Development and Evaluation for Late-Onset Alzheimer’s Disease consortium, National Institute on Aging, GSE168137, (Forner et al. 2021)).

### Quantitative histology

The brains of the transcardically perfused mice were post-perfused with 4% PFA and cryoprotected with sucrose until they lost buoyancy. They were then frozen, and sections were obtained using a cryostat and incubated with anti-V*PS*35 (clone 7E4, NBP2-78823, Novus Biologicals, CO), anti-RAB7 (clone E9O7, Cell Signaling, MA), anti-VPS13b (pa5-34406, Invitrogen) or anti-IBA1 (ab5076, Abcam) antibodies. The stacks were scanned using a Zeiss LSM700 and deconvolved. Using QuPath (Bankhead et al. 2017), we segmented cortical layer V neurons that were adjacent to the injection site but far enough from the needle track to maintain a normal parenchymal aspect. We then quantified the fluorescent signal from approximately 700 segmentations for each protein and processed and visualized the data using R. Our results are expressed as the mean +/-standard error, with confidence levels set at alpha=0.05 and two-tailed distributions used for comparisons.

## Results

### Stabilisation of the Retromer Complex by R55 in 5xFAD Mice Restores Altered Expression Of Synaptic And Neuroinflammatory Pathways

5XFAD mice were subjected to intracraneal injection by stereotax with R55 or R33, targeting an estimated final concentration of 25 μM in the parenchyma, along with vehicle-only controls. Twenty-four hours post-injection, brains were dissected for differential gene expression analysis. Out of 19,646 detected genes, 218 (1.10%) showed significant differential expression in R55-treated mice compared to vehicle controls (Figure 1A, GEO267989). R33 treatment resulted in significant changes in 19 genes (0.10%). Although most of the genes in the R55 vs. vehicle set did not reach statistical significance in the R33 vs. vehicle comparison, only 11 genes (0.05%) showed significant difference in expression between R55 vs. R33, suggesting a similar effect of the two drugs with R55 being more potent at the examined timepoint. Furthermore, only eight genes were differentially expressed in the R33 vs vehicle but not in the R55 vs vehicle gene set. Regarding the effect of the microinjection alone, 171 genes (0.87%) were significantly differentially expressed between the injected and contralateral hemisphere areas, but of these genes, only four (*Cdca5, Esyt1, Ifi30, Mmp12*) were common to the R55 vs vehicle gene set.

**Figure 1.**
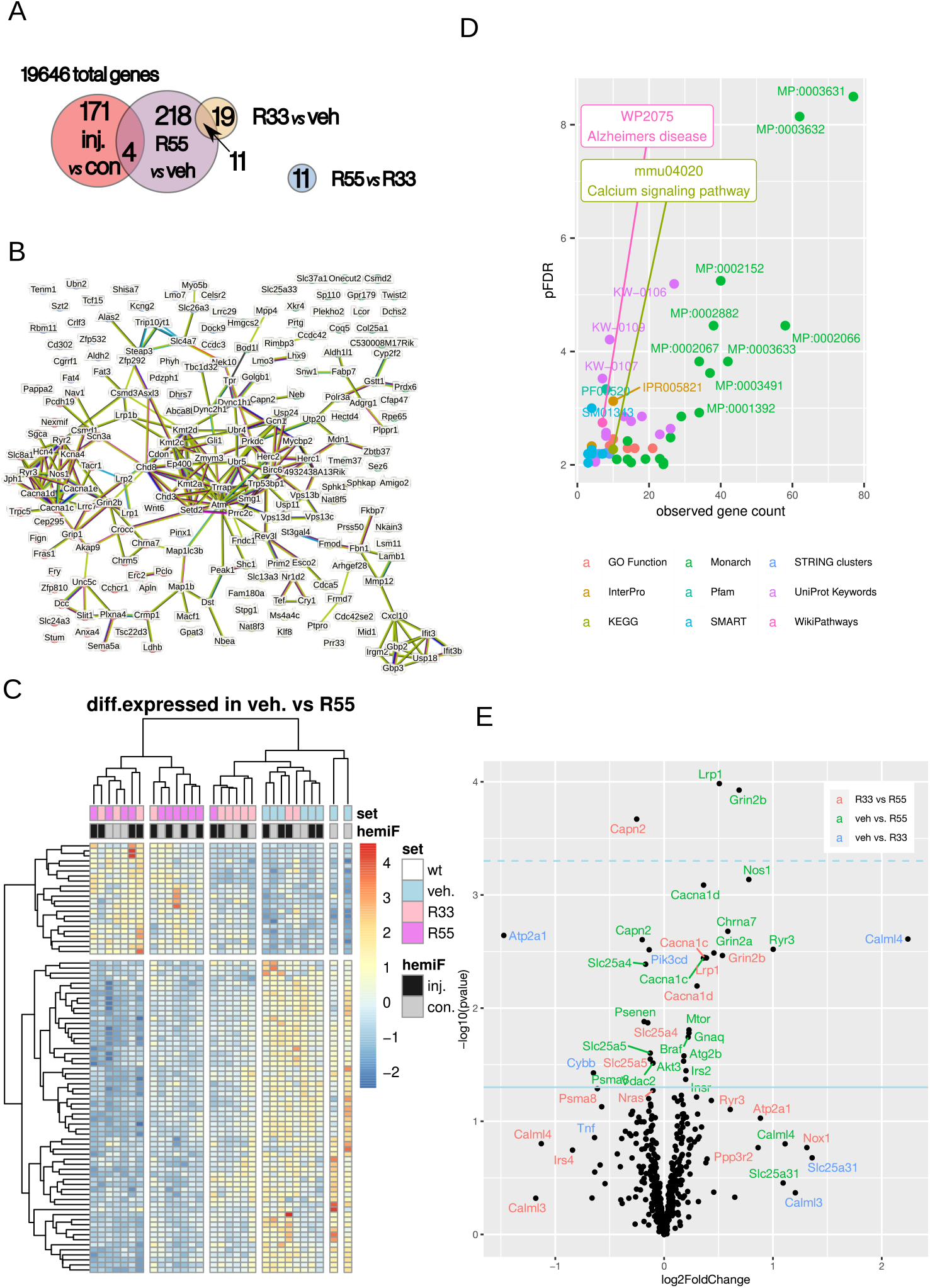
Differential expression from retromer complex stabilization by R55 and R33 in 5xFAD mice and dataset pathway enrichment. A: Summary of the number of genes differentially expressed after R55, R33, and vehicle (veh) administration in the injected (inj) and contralateral hemisphere (con). Treatment and hemispheres were included as two separate factors on the matematical model design. Constrasts were pairwise bettwen the different treatments and, separatelly, hemispheres. B: Interaction network of the DEGs between R55 and vehicle. Probability of a random genset p<1e-16. C: Heatmap of individual samples and significant hits between R55 and vehicle. D: Enrichment of the genes significantly differentially expressed between R55 and vehicle, color coded according to the database. pFDR = -log10 False Discovery Ratio. All data points shown are significant, p<0.05. E: Butterfly plot showing the significant differences in our experimental conditions vs. log2-fold changes in the expression of genes related to Alzheimer’s WP2045 dataset; the color code is based on comparisons between the 3 sets of R55, R33 and vehicle.

Network analysis of the genes differentially expressed in the R55 vs. vehicle comparison revealed 247 interactions, significantly more than the expected 101 from a random gene set (p-value = 1.10e-16), with an average node degree of 2.38 (Figure 1B). Heatmap clustering of each individual sample revealed distinct sets of gene and hemisphere pairs that were overexpressed or repressed by treatment (Figure 1C). Although R33 and R55 appeared to regulate expression at different intensities and therefore reached different statistical significance in the differential expression, the R55 and R33 samples were somehow intercalated, suggesting limited separation in their pharmacological effects. Interestingly, the microinjection of compounds influenced, as well, gene expression in the contralateral hemisphere in a similar way than the ipsilateral one suggesting widespread parenchymal distribution.Furthermore, a few individual mice injected with the drugs did not respond in the same way as the rest of the groups, suggesting some problems with drug delivery. However, due to the difficulty of objectively assigning outlier flags to these mice and because the groups were large enough to handle variations, they were not removed and included in all analysis and calculations.

Database enrichment of the significantly differentially expressed gene R55 vs vehicle set is shown in Figure 1d, which revealed a majority of the enriched neuronal and synaptic-related GO terms, as well as some related to calcium handling and axonal organization. The Alzheimer’s-related pathway WP2075 was the only significantly altered pathway, alongside significant changes in synaptic organization and other CNS-related phenotypes. Protein domain enrichment analysis highlighted domains related to PSD-95, *Egf*-like, and *Vps13* (Figure 1E).

To visualize the effect of the different comparisons, we have included a butterfly plot in Figure 1E that displays the genes of the AD related pathway WP2075. Most of the genes were changed in a similar direction by both R55 and R33. However, the effect of R55 is greater and more consistent, resulting in statistical significance. In contrast, the effect of R33 falls short of statistical significance in most cases. As R55 and R33 are highly similar structurally and the gensets R55 vs. vehicle encompass most of the R33 vs. vehicle changes, we focused mainly on R55 for the rest of the study, except on the detailed pathway illustrations where we again include both.

### Retromer Complex Stabilization Counteracts Synaptic Deficits in 5xFAD Mice

Comparison of R55-induced gene expression changes in 5xFAD mice with previously published 5xFAD vs. wild-type changes from MODEL-AD [Forner2021] revealed that R55-induced changes in the genes related to the previous enrichments were mainly of similar magnitude but in the opposite direction to those occurring in the transgenic model at 8 months, thus effectively offsetting each other. (Figure 2). Additionally, as retromer complex-related genes are not well collected in the databases, we manually curated a set of retromer genes that also included relevant cargos. To investigate if the changes relate to the AD like pathology of the 5xFAD we also compared the genes assigned to the WP2371 Parkinson’s disease related pathway, only a minority of these genes were changed, none being significant in the R55 vs. vehicle gene set. These findings indicate that R55 can compensate for many gene expression alterations associated with the 5xFAD model.

**Figure 2.**
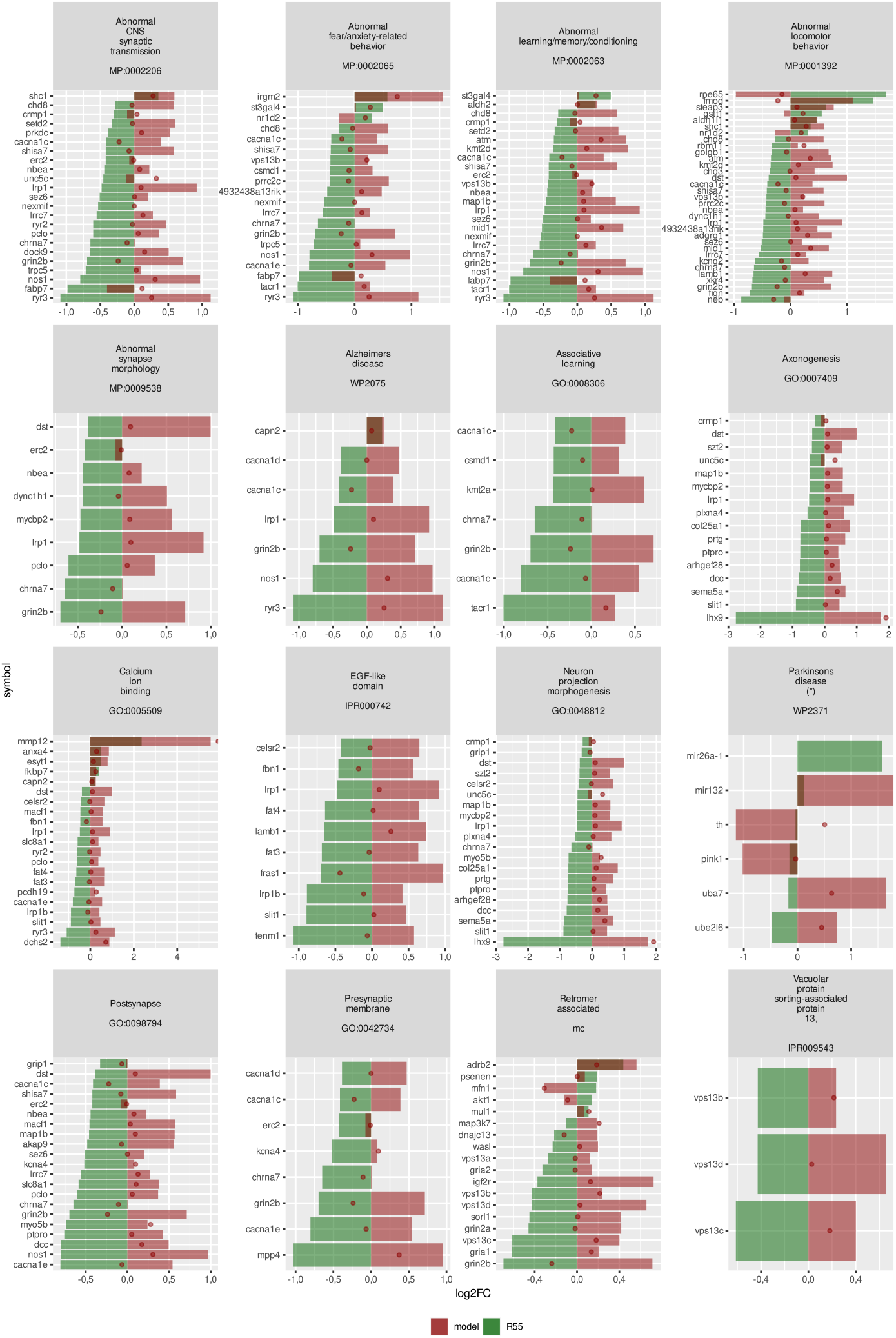
Contextualization of the pathways affected by retromer complex stabilization versus MODEL-AD. Comparison of the log2-fold changes between R55 and vehicle in 5xFAD transgenic animals (shown in green) and 5xFAD transgenic and wild-type animals of the same age (shown in red). For the latter group, there are two timepoints: dots represent 4 months, and solid bars represent 8 months from MODEL-AD dataset. Representative sets of genes were selected from different databases from enrichment analysis, and two sets were added for completion: WP2371 Parkinson’s disease and manually curated retromer-associated genes. The sets were selected to avoid overlap between them and to avoid the inclusion of large but not very informative sets, such as abnormal nervous phenotypes or abnormal brain morphology. The retromer complex-associated group was manually curated, as none of the databases that included retromer complex-associated proteins and cargos were suitable. Except for WP2371, only genes whose expression significantly changed in the R55 vs. vehicle comparison are displayed. Both genes whose expression significantly changed between the treatment group and the 5xFAD group are displayed in WP2371.

To examine alterations in synaptic-related proteins, we compared our data from R55 vs. Vehicle injection in the geneset corresponding to the abnormal learning, memory, and conditioning phenotype (monarch 2063) with previously published data on early and late long-term potentiation (LTP) [Bliim2018] and the changes in observed on the same genes in MODEL-AD. Comparing our data with the time-course data from chemically induced LTP on div15 primary hippocampal neurons at 5 minutes, 1 hour, and 2 hours post-glycine showed an increased positive correlation strength between R55-induced changes and cLTP changes as time post cLTP became longer, reaching statistical significance at 2 h (Figure 3A). Moreover, for most of these proteins, R55 treatment compensated for the 5xFAD phenotype changes, which moved in the same direction as late LTP (Figure 3B). This suggests that R55 counteracts the synaptic and LTP dysfunction observed in 5xFAD mice.

**Figure 3.**
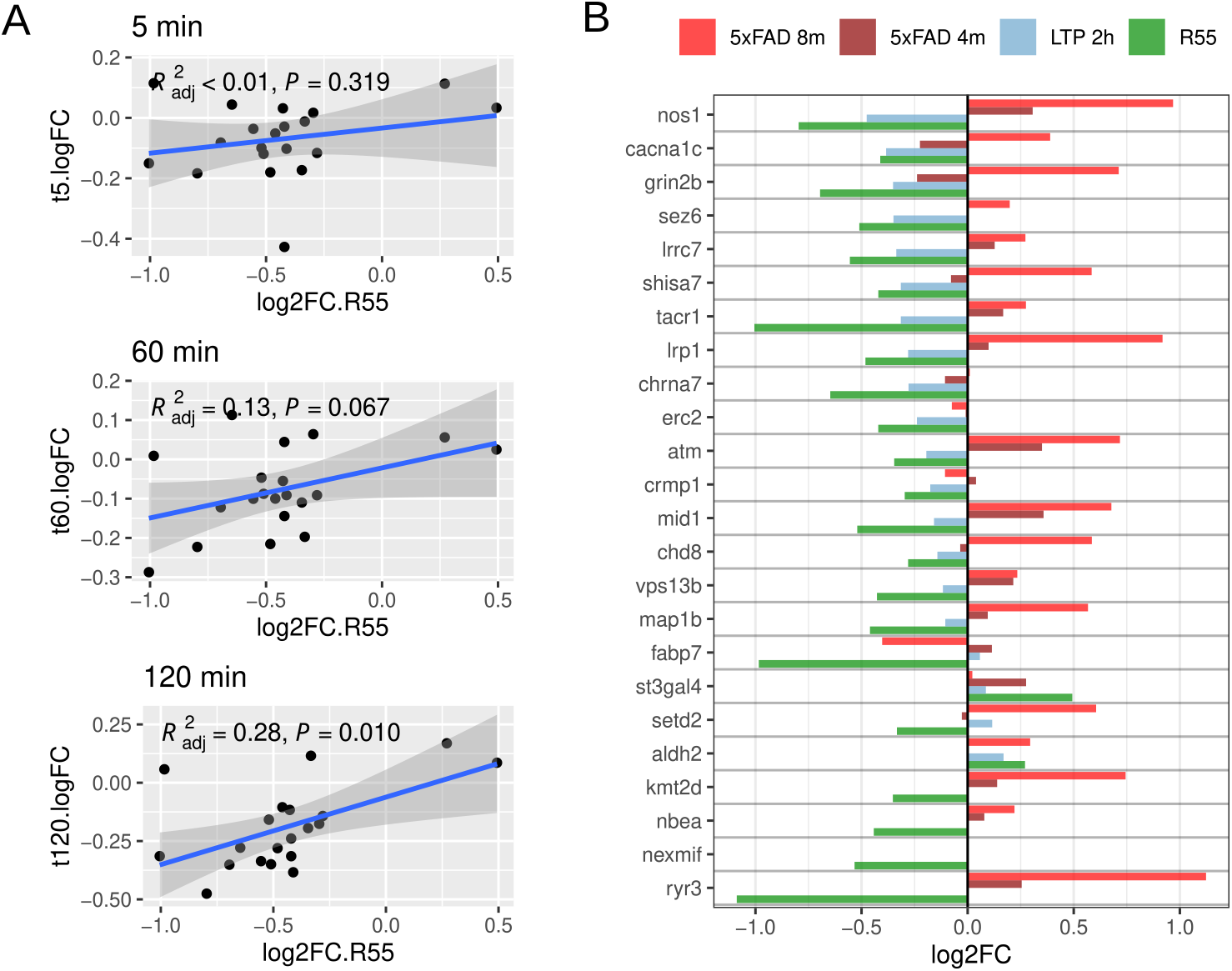
Alterations in synaptic-related proteins. A. Correlations in genes related to abnormal learning, memory and conditioning between expression changes induced by R55 (vs. Vehicle) in our mice and the changes induced at 5, 60 and 120 minutes by cLTP published by Bliim at al 2018 . B. Changes in the expression of the same genes in our dataset, in Bliim at al dataset, and in the MODEL-AD dataset at 4 and 8 months. The later ones included as reference to contextualize our findings.

In order to summarize and visualize the changes, we have mapped the expression levels induced by R33 and R55 into two annotated pathway diagrams: endosomal processing in the context of Alzheimer’s disease (Figure 4) and neuroglial interactions with a focus on the glutamatergic synapse (Figure 5). Regarding the endosome (Figure 4), it can be observed that there were no changes in the expression of the core retromer (*Vps35, Vps26a, Vps29*) or the autophagosomal-lysosomal system, except for *Lamp2* and *Pink1*. It is worth noting that although no changes in the level of expression of the core retromer complex were detected after the administration of R55, nor were they changed in the 5xFAD phenotype itself. Interestingly, however, most cargos show reduced expression, as well as consistent changes in the *Vps13* domain-containing family.

**Figure 4.**
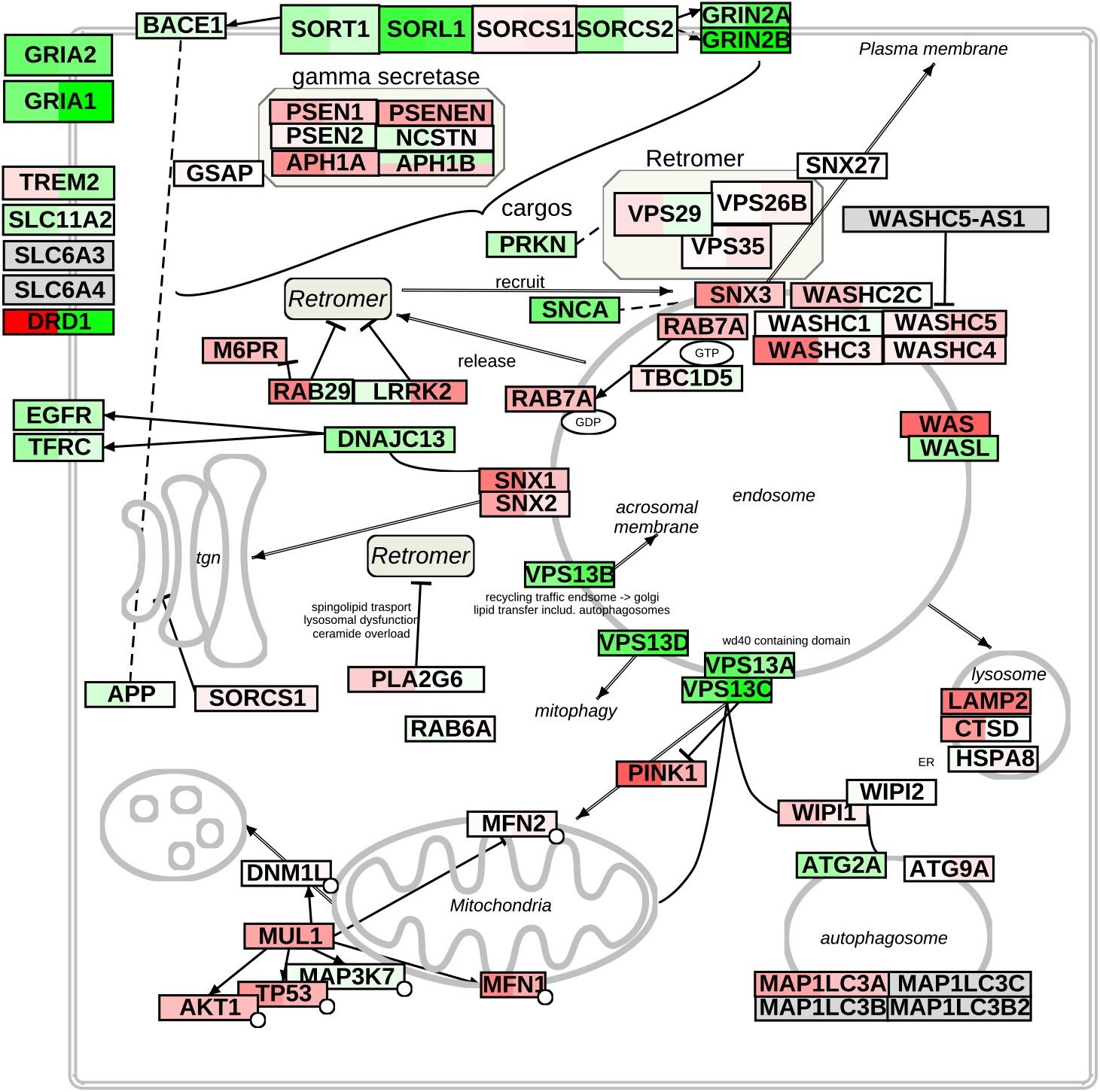
Summary of alterations in glutamatergic synapses, astrocytes and microglia over a diagram adapted from WP5083. The left box shows the changes in Z’ in our mice induced by R33, and the right box shows the changes in R55. Green indicates negative z’, and red indicates positive z’. The intensity of the tone is proportional to the degree of change. Genes with minimal and insignificant changes, which are included here for completeness, are colored almost white. Please note that this figure present the mean z-normalized values for each treatment in the injected hemisphere directly, similar to the heatmap in Figure 1C, and not log2 Fold Changes versus vehicle as in the bargraphs of figures 2, 3 and 7.

**Figure 5.**
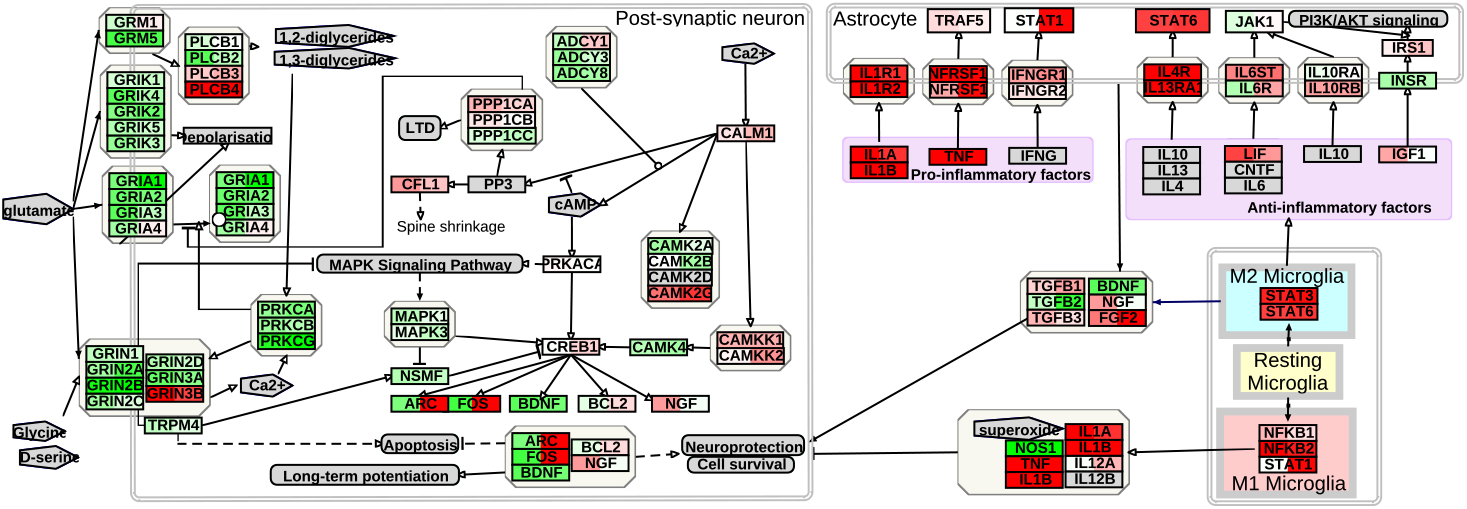
Summary of alterations in the retromer complex and related proteins relevant for Alzheimer’s disease, manually curated. The left box shows the changes in our mice in Z’ induced by R33, and the right box shows the changes in R55. Green indicates negative changes, and red indicates positive changes. The intensity of the tone is proportional to the degree of change. Genes with minimal and insignificant changes, which are included here for completeness, are colored almost white. Please note that this figure present the mean z-normalized values for each treatment in the injected hemisphere directly, similar to the heatmap in Figure 1C, and not log2 Fold Changes versus vehicle as in the bargraphs of figures 2, 3 and 7.

### Subcellular Redistribution Induced by Retromer Complex Stabilization

As alterations in the *Vps13* gene family are rather unexpected findings that could have implications for subcellular distribution, we investigated further. VPS35, VPS13b and RAB7 immunohistochemistry in the microinjected cerebral cortex after 24 hours. Interestingly, despite no significant changes in VPS35 levels were observed, concurrent with the RNA results, VPS35-positive particles were redistributed and clustered more densely and shifted away from the nuclei post-R55 treatment (Figure 6a). Consistent with the expression data, we observed a significant decrease in VPS13 and an increase in RAB7 (Figure 6b). The redistribution of VPS35 aligns with the decrease in VPS13 and the increase in RAB7 expression, suggests an enhanced or restored endosomal recycling processes.

**Figure 6.**
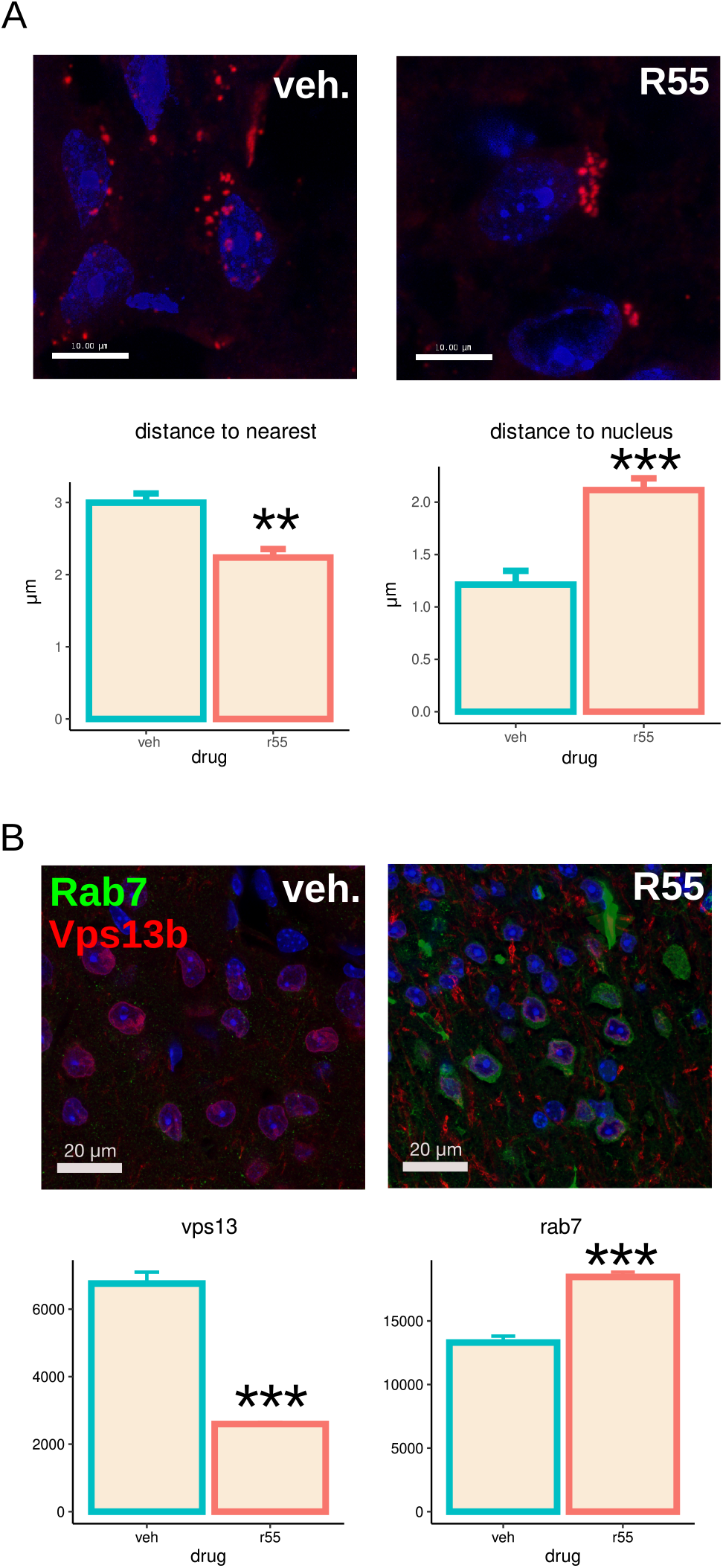
Quantitative immunocytochemistry of VPS35, VPS13 and RAB7 on the cortex of R55-or vehicle-injected 5xFAD mice. A. Representative images of the VPS35 distribution (red) after retromer complex stabilization by R55 (top) and quantification of the Vps35-positive particles (bottom). The distance to the neast partice is reduced by R55, while the distance to the nucleus increases, suggesting that the aggregation of Vps35-labeled organelles increases and the distance to the nucleus increases. B. Representative image of Vps13 (green) and Rab7 (red) after retromer stabilization by R55 (top) and quantification (bottom).

### Impact on genes related to microglial phenotype

As shown in figure 5, the treatments results in the activation of several genes responsible for microglial phenotypes, changes in astroglial receptors and the inactivation of glutamatergic synapses are observed, along with the activation of some genes downstream of Creb in neurons. *Vps35* has been described as critical for microglial activation, as it is affected by microglia-specific KO (Ren et al. 2022), so we wanted to determine the effect of stabilization of the retromer complex. We observed that IBA1-labeled microglia in R55-treated 5xFAD mice exhibited less ramification (Figure 7A), indicative of an altered activation state. As this is not easy to quantify objectively, we performed comparative analysis of gene expression changes with MODEL-AD data, this highlighted significant shifts in microglial phenotypes, including changes related to the DAM phenotype (Butovsky and Weiner 2018) and classical M0/M1/M2 subtypes (Jablonski et al. 2015) (Figures 7B and 7C).

**Figure 7.**
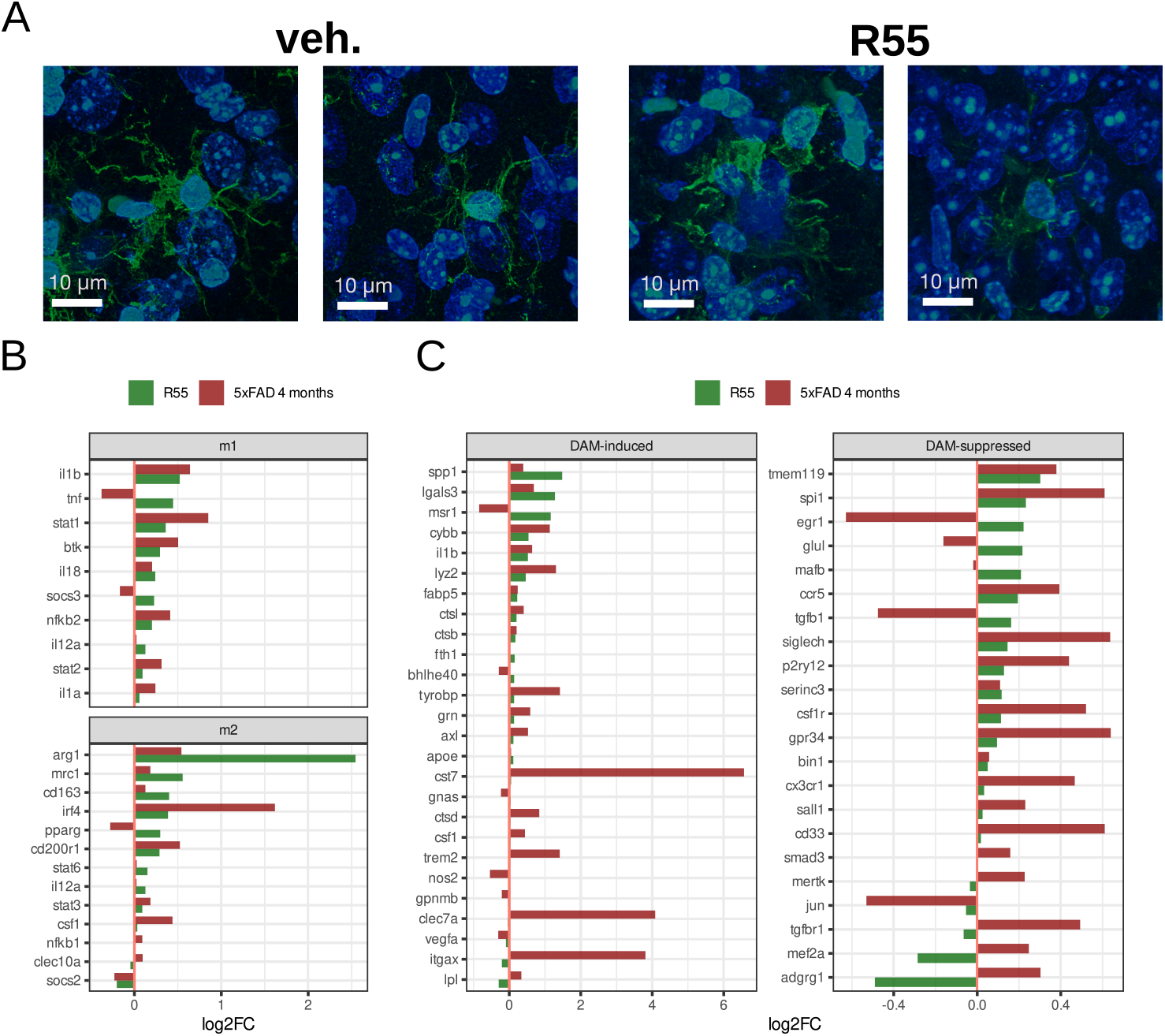
Changes in microglia after retromer complex stabilization. A. Representative images of R55-treated 5xFAD mice and vehicle-only controls. Microglia is stained with Iba-1. Microglia appears less ramified and complex after R55. B. Changes in several microglial phenotype-related genes induced by R55 in our mice contexualized with the changes in MODEL-AD at 4 months.

## Discussion

Although the exact mechanism is not yet fully elucidated, a robust association exists in the literature between AD progression and synaptic dysfunction. It is believed that in the early stages, Aβ oligomers directly interfere and induce reversible changes similar to those involved in developmental synaptic pruning and normal plasticity, such as LTP and LTD. These phenomena have been extensively studied in vitro (S. Li and Stern 2022). Progressively, it appears that interactions between Aβ and different proteins lead to irreversible synaptic changes. For instance, toxic Aβ oligomers have been shown to increase cholesterol concentration at synapses, promoting Aβ/PrP complexes that decrease NMDAR affinity for glycine and ultimately lead to the inactivation of GRIN2a and GRIN2b subunits via S-nitrosylation (Benarroch 2018). These events are reflected in the MODEL-AD dataset for 5xFAD mice as apparent compensatory increases in the expression of *Grin2a*/*Grin2b, iNos*, and certain VLDL/ApoE-related genes such as *Lrp1, Lpr1b*, and *Fbn1*. Notably, in our study, retromer complex stabilization by R55 significantly counteracted these elevated levels, likely due to the stabilisation of the retromer enhance their membrane presence, thereby reducing the need for their upregulation.

The retromer complex has been implicated in proper synaptic plasticity, particularly involving *Gria1 (*AMPA) receptors. Studies have demonstrated that shRNA-mediated silencing of *Vps35* in hippocampal cultures not only disrupts AMPA receptor membrane insertion but also impairs LTP, whereas restoring *Vps35* reverses these effects (Temkin et al. 2017). In our study, administering R55 at an age known for impaired LTP effectively counterbalanced the altered expressions of *Gria1* and *Grip1*. This subtle compensation of *Grip1* is notable, as it is pivotal for the endosomal sorting essential for AMPA receptor recycling and membrane targeting. Recent research on AMPA and LTP dynamics have implicated the semaphorin family, specifically Plexin a4 *(Plxna4a)*, in driving AMPA receptors to synapses and facilitating LTP-associated plasticity (Jitsuki‐Takahashi et al. 2021). In the 5xFAD model, the expressions of *Plexin a4*, semaphorin 5a (S*ema5a*), and the netrin receptors *Unc5c* and *Dcc* are altered, which we found to be compensated by retromer complex stabilization.

Disrupted calcium handling, particularly intracellular calcium leakage from the endoplasmic reticulum (ER), has been implicated in early synaptic dysfunction observed in Alzheimer’s disease. Alterations in neuronal *Ryr2* and changes in ryanodine binding have been detected from early stages of the disease and throughout its progression, resulting in both enhanced receptor-operated calcium entry and decreased buffering capabilities (Kelliher et al., n.d.; Lacampagne et al. 2017). Furthermore, ryanodine receptors (RyRs) exhibit functional coupling with L-type calcium channels, particularly those involved in calcium-induced calcium release (CICR), especially in CA1 pyramidal neurons (Sahu and Turner 2021). Our findings show that retromer complex stabilization compensated for the alterations observed in the expression of *Ryr2-* and Ryr2-interacting L-type calcium channels (*Cacna1c, Cacna1d*, and *Cacna1e*). Additionally, of the known interactors of *Ryr2* that regulate its hyperexcitability (Takeshima, Hoshijima, and Song 2015), such as *Fkbp1b, Mapk14, Mapk3*, and *Jph3/4*, only *Jph4* is weakly modified by R55 We speculate that normalizing retromer function could restore early endosomal recycling, thereby obviating the need for pathological compensation.

Concerning AD-related proteins and known retromer complex cargos, R55 treatment increased the levels of the Aβ-processing gamma-secretase components and notably reduced the levels of *Sorl1* and *Col25a1*, the latter a collagen-like amyloid plaque component known for producing protease-resistant aggregates. This could indicate that excessive processing by presenilins, due to extended cargo residency at the Golgi in dysfunction, might account for the observed compensatory effects. However, unexpectedly, R55 treatment also led to a reduction in all members of the *Vps13* family that were increased in the Alzheimer’s disease model. *Vps13* directs recycling endosomes towards the Golgi, mitochondria, or autophagosome vesicles by promoting organelle bilayer bridging. Notably, *Vps13c* have been identified in GWAS as early-onset atypical PARK23 and related to mitochondrial clearance deficiencies in neurons, a known hallmark of Parkinson’s disease (Jitsuki‐Takahashi et al. 2021; Ugur et al. 2020). Genetic variation in VPS13 has been identified in a large-scale multivariate modelling of genotype-phenotype structural MRI of sporadic, late onset AD, with risk contributions exceeding even ApoE (Meda et al. 2012). Immunohistochemistry confirmed a decrease in neuronal Vps13 protein levels and an increase in Rab7, consistent with our gene expression findings, suggesting restored endosomal trafficking to the plasma membrane following retromer stabilization. This also explains why Vps35 labeling in R55-treated mice was distributed farther from the nucleus compared to that in vehicle-treated mice, indicative of reduced Golgi association.

In the neuroinflammatory unit, R55 treatment not only increased the levels of pro-inflammatory mediators (such as *Il-1* or *Nfkb2*) but also enhanced the levels of pro-trophic glial markers (*Il4r* and *Il13ra1*) and downstream anti-inflammatory transcription factors (*Stat6* and *Stat3*). This coordinated alteration in crosstalk between microglial polarization and astrocytic responses involving *Il1, Il4, Il13, Nfkb*, and *Stat6* mirrors findings reported in other AD studies (Xie et al. 2020). R55 also influenced the expression of microglial genes involved in the DAM phenotype, as well as in the former M0/M1/M2 model, as indicated by more than 2-fold changes in the expression of *Arg1*. These findings are opposite too and consistent with previous reports where *Vps35* KO impaired microglial activation in response to Aβ, affecting both pro-inflammatory and trophic phenotypes (Ren et al. 2022). As phagocytic receptors, such as cytokine receptors, share the same pathway from early endosomes to the membrane in glial cells as the synaptic receptors in neurons, the restoration of retromer complex function by R55 offers a plausible explanation for these observed changes.

While extrapolating from acute to long-term chronic protection remains challenging, focusing on the pathophysiological mechanism, our data support the involvement of retromer dysfunction in AD and suggest that early synaptic dysfunction could be mitigated by retromer stabilization strategies. It remains an open question why retromer dysfunction results in different outcomes, leading to Parkinson’s and Alzheimer’s diseases, but perhaps the effects of two different VPS35 mutations, D620N and L625P, associated with each disease respectively, could provide some understanding. The Parkinson’s-related D620N mutation reduces binding to the actin nucleation complex, *Wash*, which is unaltered in AD models (MODEL-AD) and unaffected by R55 (Berman et al. 2015), while the Alzheimer’s-related L625P mutation directly impacts VPS35 assembly with VPS26 and VPS29 (McGough et al. 2014; Rovelet-Lecrux et al. 2015). This could highlight an essential role for retromer anchoring, ultimately determining subcellular localization and microdomains. The ability of R55 to rapidly influence retromer complex distribution and the expression of membrane-related anchors such as VPS13 supports this hypothesis.

In summary, we have identified a number of retromer cargos, which are membrane-associated proteins, receptors, or part of the post-translational processing machinery, that are altered in early AD and are rapidly compensated for by retromer stabilization. Those are involved in the pathophysiological mechanisms of microglial activation and early synaptic dysfunction in AD, particularly around the maintenance of long-term potentiation. We have also shown that retromer stabilization involves a cellular redistribution of VPS35-containing particles to a position further away from the nucleus and that this is accompanied by changes observed in late endosomal to Golgi anchoring components such as VPS13 and late endosomal recycling to the membrane such as RAB7, providing a coherent interpretation of the redistribution as more efficient recycling. We believe this is relevant to the exploration of VPS35 stabilization as a potential therapeutic strategy for AD, and as this work was performed in 5xFAD mice, which do not carry genetic alterations in VPS35 or any retromer component, this strategy may also be generalizable to the more common sporadic forms of the disease.

## Declarations

### Ethics approval and consent to participate

All procedures and working protocols were approved by the animal ethical committee of Linköping University.

## Consent for publication

no applicable as does not contain any data from individual persons.

## Availability of data and material

Datasets have been deposited at GEO267989. Raw images and histological data are avaible upon request to the data curators.

## Author contributions

DR, MB: designed research and conceived the study. DR, AN: performed experimental work. DR: developed or adapted methodology. DR: performed computational work. DR, MB: collected and are curating the data. DR: analysed and interpreted the data. DR: wrote the manuscript. MB: provided funding and physical framework. All authors reviewed the manuscript.

## Acknowledgements

The authors express gratitude to Åsa Schippert from the Molecular Biology Unit of the LiU’s Core Facility for her assistance with RNA deep sequencing, Vesa Loitto from the Microscopy Unit of the LiU’s Core Facility for his assistance with imaging, and Sarah Lindström for her genotyping and maintenance of the 5xFAD mice colony and her assistance in the Animal Facility.

